# Intracellular acidification by bacteria-derived valeric acid is a mechanism of trans-kingdom ecology against *Candida parapsilosis* colonization

**DOI:** 10.1101/2025.09.12.675697

**Authors:** Keiko Yasuma-Mitobe, Chen Liao, Tibor Nemeth, Kevin Byrne, Audrey Bilipps, Ruben Jose Jesus Faustino Ramos, Cristine Nicole Salinas, Eric Chan, Martina Perissinoto, Ashley M. Sidebottom, George Plitas, Geraldine Butler, Justin R Cross, Eric G Pamer, Attila Gacser, Joao B Xavier, Tobias M. Hohl

## Abstract

In hematopoietic cell transplant patients, intestinal *Candida parapsilosis* expansion and translocation cause life-threatening candidemia, yet how commensal intestinal bacteria prevent *Candida* expansion remains incompletely defined. Here, we trained a machine learning model on supernatant metabolomic profiles of commensal bacteria to identify bacteria-derived inhibitors of fungal growth, with valeric and butyric acid as top hits. Experimental validation confirmed *in silico* predictions in three systems. First, in patient fecal samples, valeric and butyric acid levels inversely correlated with *C. parapsilosis* growth. Second, *in* vitro, valeric acid potently inhibited *C. parapsilosis* growth by causing intracellular acidification. Third, administration of glycerol valerate, and free or microencapsulated valeric acid blunted *C. parapsilosis* growth at murine intestinal sites where valeric acid could be detected. Thus, machine learning could identify a mechanistic driver of trans-kingdom ecology limiting *C. parapsilosis* intestinal expansion and may inform strategies to reduce patient risk of developing candidiasis during high-risk periods.

## Introduction

*Candida* species are members of the intestinal microbial ecosystem and asymptomatically colonize ∼50-70% of individuals, with expansion checked by bacterial competitors and host immune effectors^1, 2^. However, *Candida* species, including *C. albicans*, *C. parapsilosis*, *C. glabrata*, and *C. auris,* can invade mucosal barriers and cause life-threatening bloodstream infections (BSIs), primarily in critically ill patients, in patients with hematologic malignancies, and in solid organ and hematopoietic cell transplant (HCT) recipient^3, 4, 5, 6, 7, 8, 9, 10, 11, 12, 13^.

In these high-risk groups, patients often receive antibiotics to prevent or treat bacterial infections, injuring the commensal bacterial ecosystem and facilitating the expansion of *Candida* species and other pathobionts. For example, longitudinal analysis revealed that ∼20-30% of allogeneic HCT (allo-HCT) patients exhibited *Candida* intestinal expansion, and that a subset of these patients developed systemic candidiasis despite concurrent receipt of antifungal prophylaxis^14, 15^. Moreover, *C. parapsilosis* and *C. albicans* intestinal expansion have been linked to adverse HCT transplant outcomes, including non-relapse mortality^14, 16^

Analysis of intestinal bacterial populations showed that the loss of *Lachnospiraceae*^15^ and the presence of lactate-producing species, e.g., *Lactobacillus*, *Lactococcus, Klebsiella*, and *Escherichia,* correlated with *Candida* abundance^17^. These observations and predictive modeling advanced the concept that commensal bacteria engage in trans-kingdom ecology with fungi, namely, through bacteria-derived metabolites that regulate *Candida* colonization levels. Studies in murine models indicated that commensal bacteria could trigger hypoxia-inducible factor 1α-dependent secretion of cathelidin-related antimicrobial peptide (CRAMP; LL-37 in humans) to limit *C. albicans* expansion^18^. Host production of *C. albicans*-specific IgA and antimicrobial peptide YY can preferentially target the *C. albicans* hyphal morphotype that mediates tissue invasion and virulence^19, 20, 21, 22, 23^. Moreover, commensal bacteria contribute to hypoxic environments in the intestinal tract, and pharmacologic strategies that promote anaerobiosis limit *C. albicans* colonization^24^. *In vitro* studies support the idea that 2-, 3-, and 4-carbon short-chain fatty acids (SCFAs) and bile acids may inhibit *Candida* growth^20, 25, 26, 27, 28, 29, 30, 31^, including *C. albicans* hyphal formation and cell wall remodeling^32, 33, 34^. However, it remains unclear whether changes in SCFA or other bacterial metabolites could mediate trans-kingdom ecology in the intestinal tract, and if so, what the underlying mechanisms of *Candida* growth inhibition are.

In this study, we trained a machine learning model using the secreted metabolomes from commensal bacteria as an input to identify bacterial-derived valeric acid as a potent and direct inhibitor of *C. parapsilosis* growth. This prediction was validated through correlative analysis of valeric acid abundance and *C. parapsilosis* growth in human fecal specimens, through *in vitro* studies that revealed valeric acid-mediated loss of *C. parapsilosis* intracellular pH homeostasis and growth inhibition. Delivery of valeric acid formulations demonstrated *C. parapsilosis* growth inhibition at gastrointestinal sites where valeric acid could be detected. These results highlight the potential of harnessing machine learning models to identify and characterize bacterial-derived metabolites as a novel strategy to limit *Candida* colonization and mitigate risk for life-threatening infections.

## Results

### Machine learning identifies bacterial-derived valeric acid and other SCFAs as candidate inhibitors of *Candida* growth

To quantify the effects of secreted bacterial metabolites on *C. parapsilosis* growth, we cultured a clinical *C. parapsilosis* isolate (strain 1004) in a 1:1 mixture of fresh brain heart infusion (BHI) broth and spent BHI supernatant of 347 human commensal bacteria, primarily *Lachnospiraceae*^35, 36^. To account for nutrient depletion, control cultures contained a 1:1 mix of fresh BHI and phosphate-buffered saline (PBS; Figure 1a). Spent supernatants from *Coprococcus comes*, *Anaerostipes hadrus,* and *Blautia wexlerae* isolates inhibited *C. parapsilosis* growth, as judged by the area under the curve, compared to control growth conditions (Figure 1b). Notably, *C. comes* and *A. hadrus* are known SCFA producers^37, 38^.

**Figure 1.**
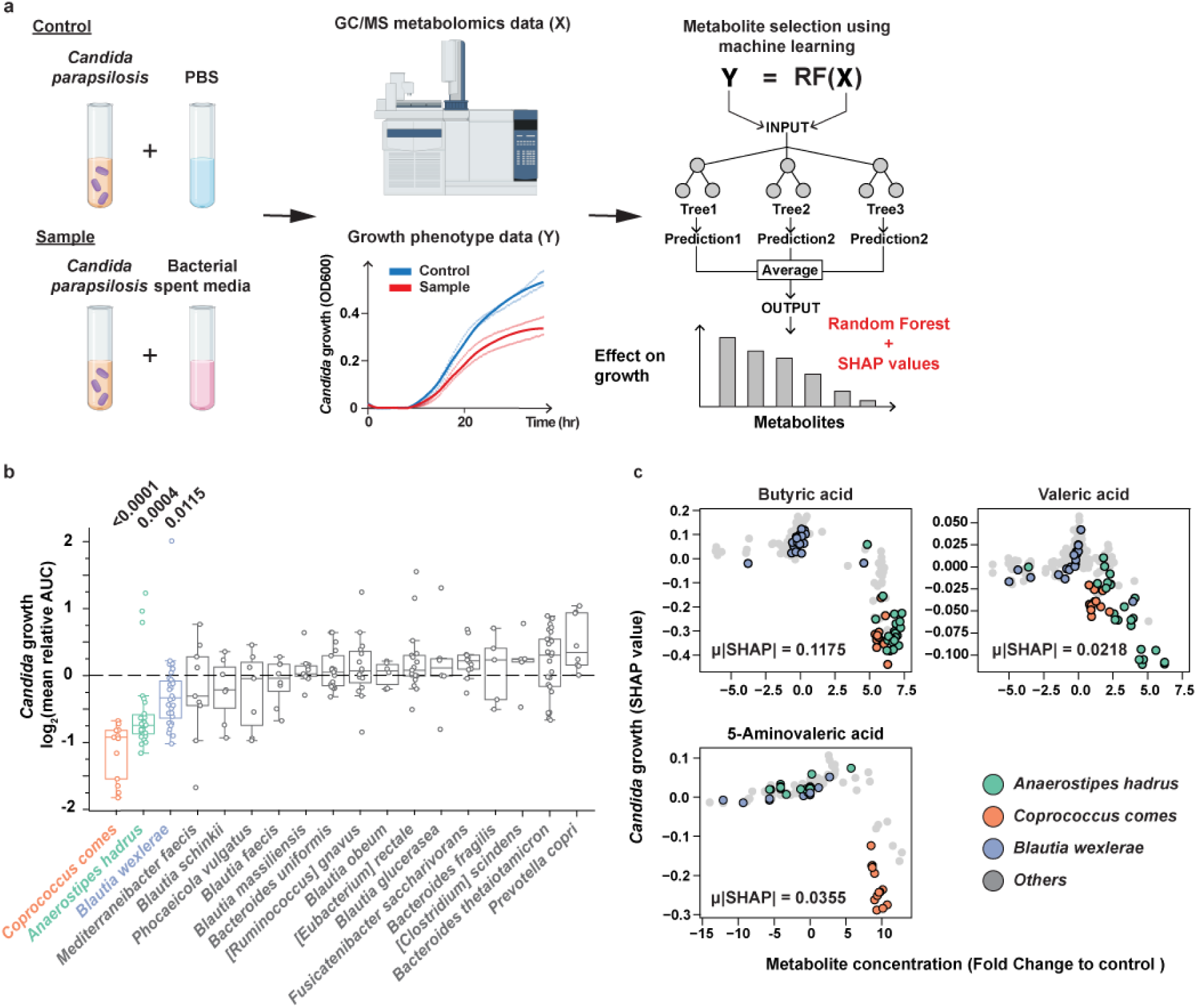
Prediction of bacterial metabolites that inhibit *C. parapsilosis* growth. **a,** Schematic summary of screen, analysis, and mathematical modeling to identify *C. parapsilosis* growth inhibitory metabolites from spent bacterial supernatants. *C. parapsilosis* strain 1004 was resuspended in BHI broth and added to an equal volume of spent bacterial supernatant (Sample) or PBS (Control) and growth was measured by OD_600_ area under the curve (AUC) analysis at 24 h. Spent bacterial supernatants were analyzed by GC-MS to measure the abundance of 36 targeted metabolites to develop a Random Forest model and predict *Candida* growth from the metabolomic composition of spent supernatants. SHAP analysis calculated the impact of individual candidate metabolites on *C. parapsilosis* growth. **b**, *Candida* AUC ratio in individual spent bacterial supernatants compared to control samples shown in log_2_ scale. Isolates are shown by species, with only species with 5 or more isolates displayed, ordered by median AUC ratio. Unclassified isolates are excluded. **c**, SHAP analysis of *Candida* growth AUCs (y axis) in relation to the concentration of butyric, valeric, and 5-aminovaleric acid (x axis). Each dot represents one bacterial isolate. *Coprococcus comes*, *Anaerostipes hadrus*, and *Blautia wexlerae* isolates are highlighted in orange, green, or blue as shown. The means of the absolute SHAP values (μ|SHAP|) are shown. Statistical analysis: **b**, One sample t-test with Benjamin-Hochberg correction; adjusted *P* values are shown.

To identify candidate inhibitory metabolites, we compared the abundance of 36 target metabolites in 229 spent supernatants with their abundance in fresh BHI broth (Figure 1a, Extended Figure 1, Supplementary Table 1). We used these metabolomics data to fit a machine learning model of *Candida* growth. Then, we used Shapley a*dditive exPlanations* (SHAP) analysis on a Random Forest model to determine the impact of each bacteria-derived metabolite. This model showed good prediction performance with an R² of 0.75, indicating *Candida* growth could be explained by the metabolite features (Extended Figure 2a). The model identified valeric acid, 5-aminovaleric acid, and butyric acid as inhibitors of *Candida* growth, as well as other SCFAs and amino acids (Figure 1c, Extended Figure 2b). These candidate inhibitors were first investigated in human allo-HCT patient fecal samples.

### Valeric acid and other SCFA abundances inversely correlate with *Candida parapsilosis* colonization in hematopoietic cell transplant patient fecal samples

To corroborate predicted metabolomic hits with *C. parapsilosis* intestinal colonization data, we analyzed fecal samples from 22 allo-HCT patients (145 fecal samples, range: 2-17 per patient, Supplementary Table 2 for profiles) that were on micafungin prophylaxis (see Materials and Methods). Each allo-HCT recipient had both fungal culture-positive (n = 64) and fungal culture-negative fecal samples (n = 81). For each sample, we analyzed 43 metabolites by gas chromatography–mass spectrometry (GC-MS), including the abundance of 2- to 5-carbon SCFAs and 5-aminovaleric acid. In addition, the relative abundance of bacterial and fungal taxa was analyzed by 16S and ITS1 rDNA sequencing, respectively, and the *C. parapsilosis* colony-forming units (CFUs) were enumerated.

A patient example illustrates a temporal relationship between fecal SCFA abundances and *C. parapsilosis* colonization (Figure 2a). The patient experienced loss of *Lachnospiraceae* starting on day −4 of allo-HCT, which preceded a >4 x log_10_ reduction in the peak area of valeric acid and other SCFAs on day −1 of allo-HCT. These events occurred prior to *C. parapsilosis* expansion as judged by the rise in relative *C. parapsilosis* ITS1 rDNA abundance and the recovery of viable *C. parapsilosis* CFU on day +1. After the patient experienced engraftment on day +11 and recovery of SCFA fecal abundances on day +16, *C. parapsilosis* ITS1 expansion declined and the fungus was undetectable by culture. Among all samples tested, fecal valeric acid, 5-amino valeric acid, butyric acid, propionic acid, and acetic acid peak values negatively correlated with fecal fungal CFU (Figure 2b-2f; Extended Figure 3). Samples with high peak SCFA abundances were nearly uniformly *C. parapsilosis* culture-negative, consistent with the idea that valeric acid and other 3-5 carbon SCFAs inhibit *C. parapsilosis* expansion in the human intestinal tract.

**Figure 2.**
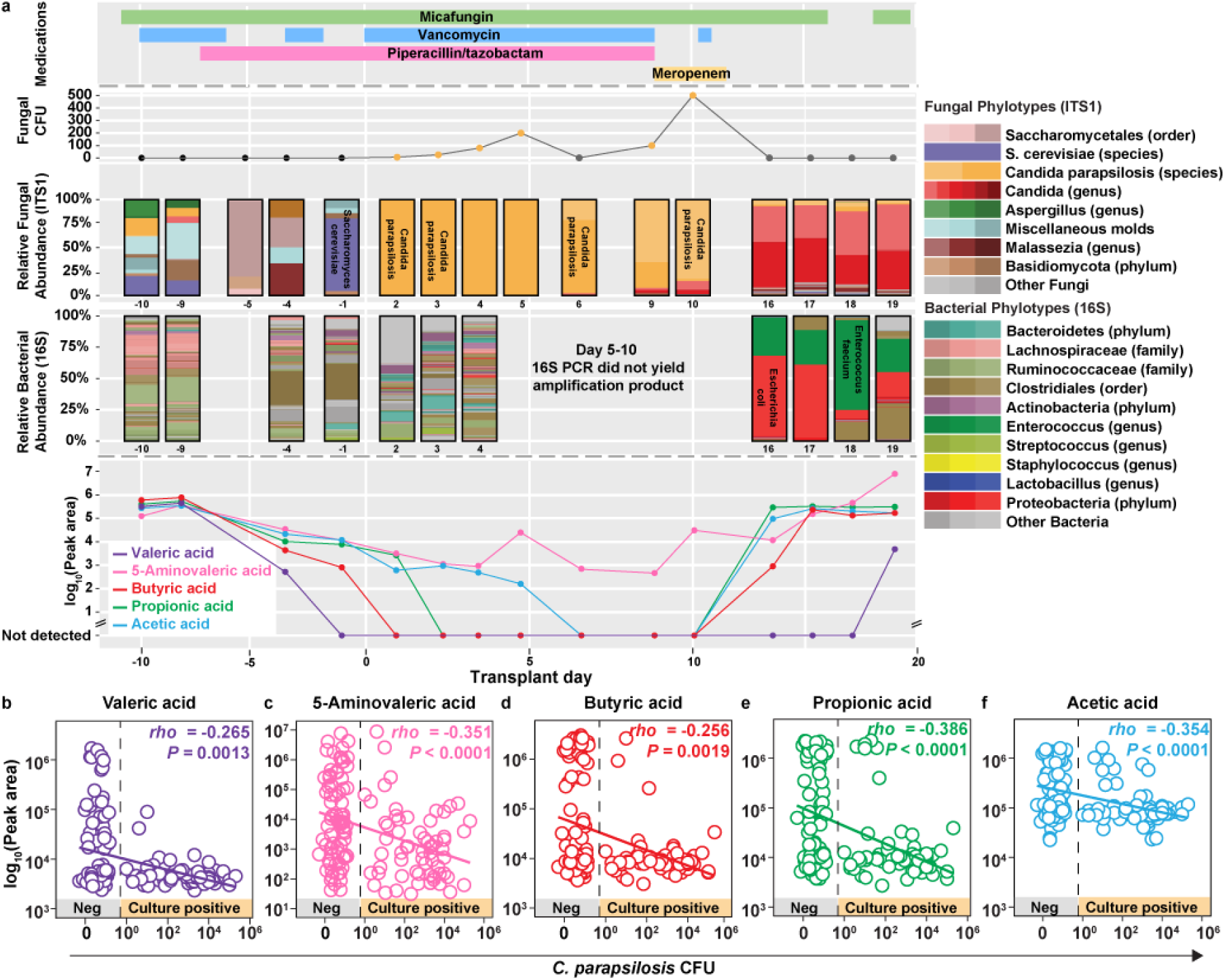
Allo-HCT patient fecal profiles for *C. parapsilosis* and candidate metabolites. **a**, Longitudinal profile of one allo-HCT patient from day −10 to day +20, with rows indicating antimicrobial medication (top), intestinal fungal colonization by fecal CFU (second), relative abundance of fungal taxa by ITS1 amplicons (third), relative abundance of bacterial taxa by 16S amplicons (fourth), and the abundance of indicated metabolites by GC-MS peak area (bottom). **b-f**, The plots indicate *C. parapsilosis* CFU and (**b**) valeric acid, (**c**) 5-aminovaleric acid, (**d**) butyric acid, (**e**) propionic acid and (**f**) acetic acid peak areas in each sample (n = 145 fecal samples from 22 allo-HCT patients). Statistical analysis: **b-f,** Spearman’s *rho* and two-tailed *P* values are shown.

### Valeric acid inhibits *Candida* growth in vitro

To evaluate *in silico* predictions, we tested the effect of four SCFAs (i.e., valeric, butyric, propionic, and acetic acid) and 5-aminovaleric acid on the growth of *C. parapsilosis* and other *Candida* species. At 6 mM concentration, valeric acid inhibited *C. parapsilosis* growth in a more potent manner than butyric acid and propionic acid, while acetic acid did not inhibit growth (Figure 3a-3b). Similar results were observed with *C. metapsilosis* and *C. albicans* (Extended Figure 4a-4b). In contrast, 5-aminovaleric acid did not inhibit C. *parapsilosis* growth (Figure 3c), suggesting that addition of an amino group to valeric acid resulted in loss of antifungal activity.

**Figure 3.**
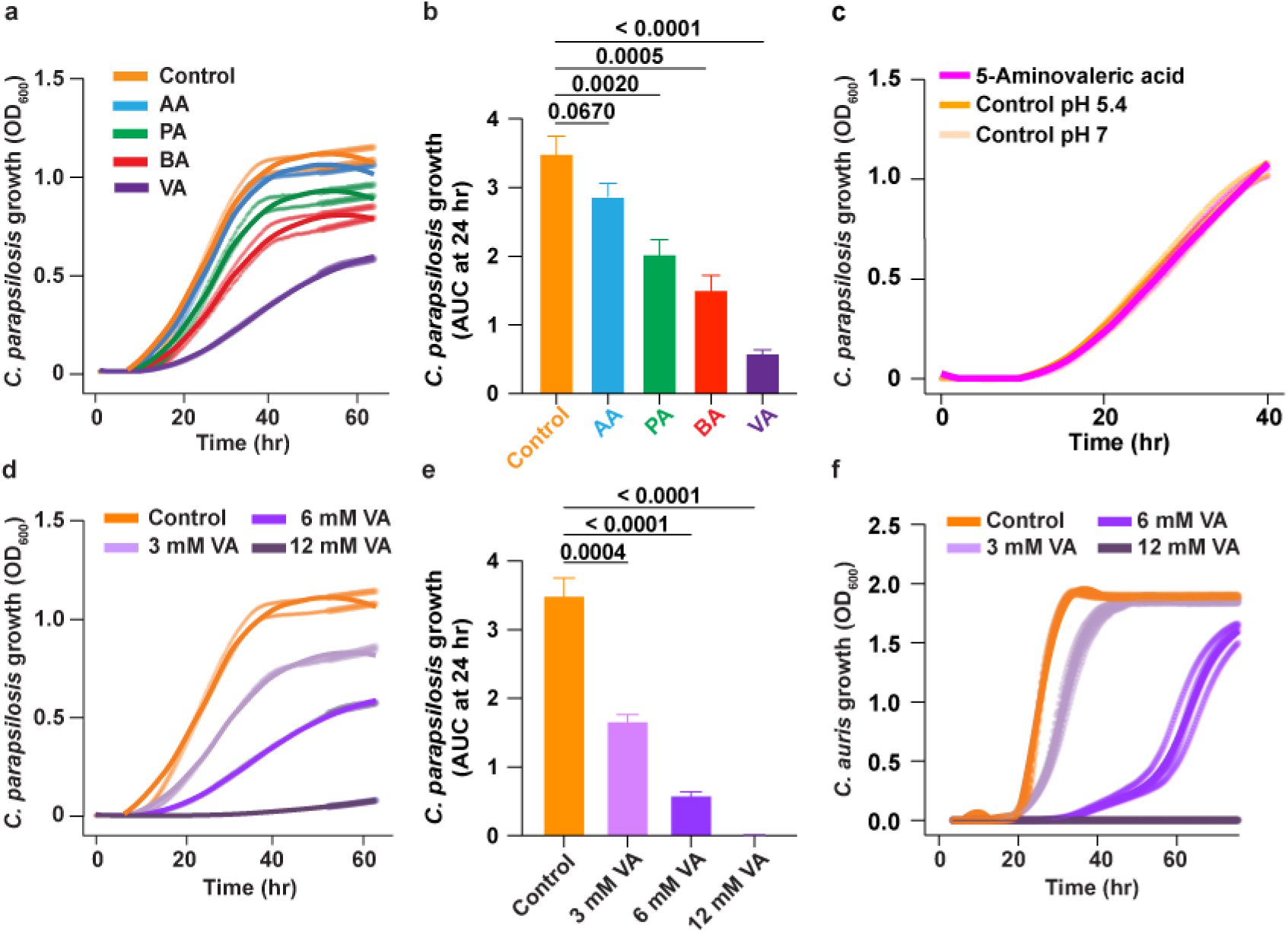
Effects of short-chain fatty acids on *Candida* growth. **(a, c, d)** *C. parapsilosis* 1004 or (**f**) *C. auris* 1193 growth kinetics by optical density (600 nm) and (**b, e**) *C. parapsilosis* AUC values at 24 hr after culture in YPD broth with (**a, b**) 0 mM (control, orange symbols) or 6 mM of the indicated SCFAs (AA, acetic acid, light blue symbols; PA, propionic acid, green symbols; BA, butyric acid, red symbols; VA, valeric acid, purple symbols), with (**c**) 0 mM (pH 5.4, orange line or pH 7.0, yellow line; both adjusted with HCl) or 12 mM 5-aminovaleric acid (pink line), with (**d-f**) 0 mM (control, orange symbols), 3 mM (light purple symbols), 6 mM (purple symbols), or 12 mM (dark purple symbols) valeric acid. Data are representative of at least two independent experiments and growth curves were calculated from at least two replicates in an experiment. Statistical analysis: **b, e,** Adjusted *P* values versus control, calculated with one-way ANOVA with Dunnett’s multiple comparison test.

**Figure 4.**
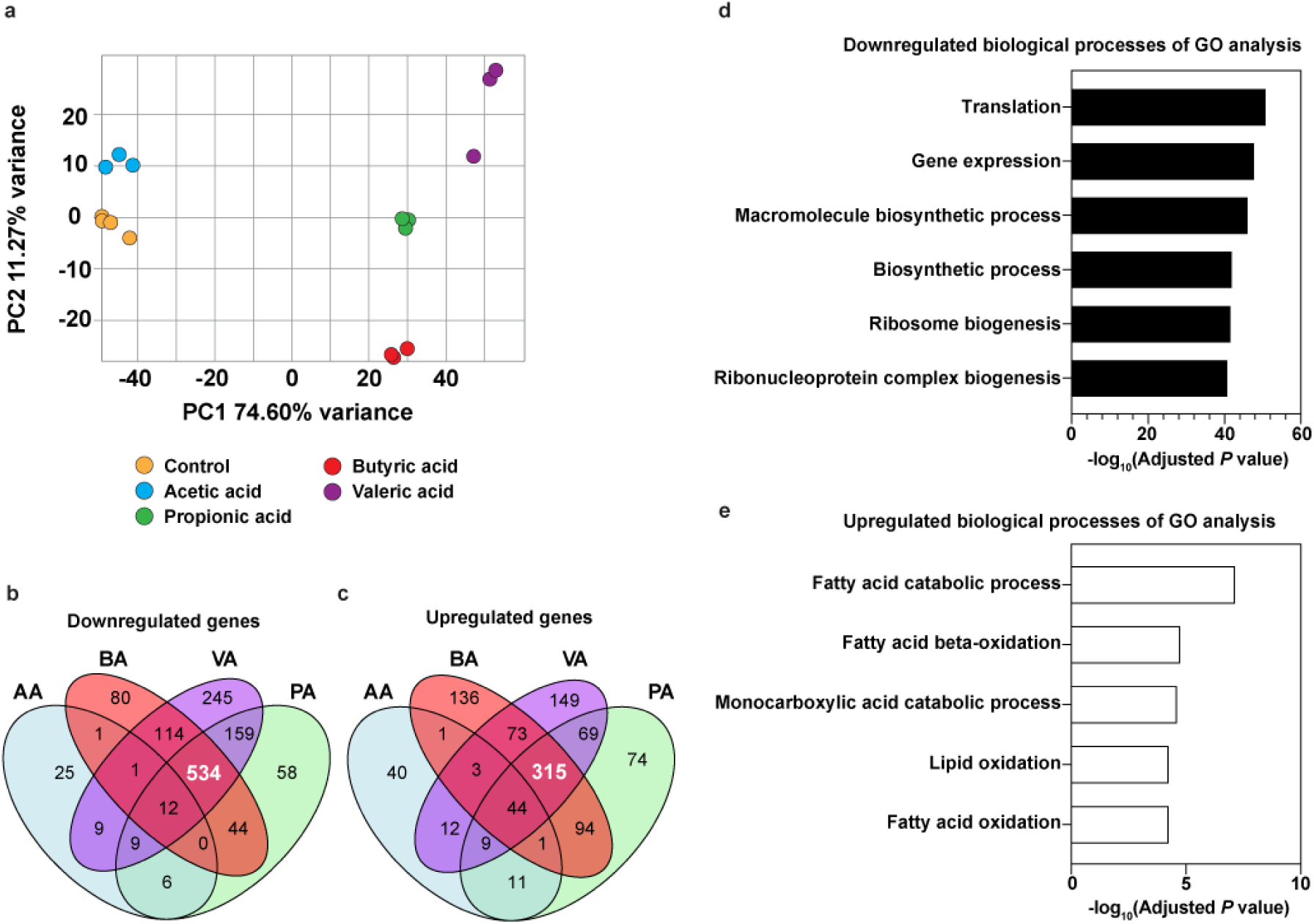
Transcriptome analysis of SCFA-treated *C. parapsilosis* cells.

Valeric acid-mediated growth inhibition occurred in a dose-dependent manner (Figure 3d-3e) and across all tested *C. parapsilosis* (Figure 3e and Extended Figure 4c-4d), *C. orthopsilosis* (Extended Figure 4e-4f), *C. metapsilosis* (Extended Figure 4g-4i), *C. albicans* (Extended Figure 4j-4l), *C. glabrata* (Extended Figure 4m), *C. tropicalis* (Extended Figure 4n), and *C. auris* (Figure 3f) isolates. Thus, valeric acid exposure inhibits *Candida* growth across multiple species.

### Valeric acid exposure alters *C. parapsilosis* gene expression

To understand how bacteria-produced SCFAs may be inhibiting *C. parapsilosis* growth, we performed transcriptome analysis in the presence of 0 or 25 mM valeric, butyric, propionic, or acetic acid. Principle component analysis (PCA) revealed a minor difference in gene expression between the control and the acetic acid samples, and a large difference between these and propionic, butyric, or valeric acid-treated samples (Figure 4a), consistent with the finding that acetic acid did not inhibit *Candida* growth (Figure 3a-3b, Extended Figure 4a-4b). Overall, 63, 822, 786, and 1,083 genes were downregulated by acetic, propionic, butyric, and valeric acid treatment compared with the control condition, representing a total 1,297 unique genes. Of these, 534 downregulated genes were common to valeric, butyric, and propionic acid exposure (Figure 4b, Supplementary Table 3). Gene ontology (GO) analysis of these 534 genes revealed that pathways related to ribosomal function and translation were downregulated (Figure 4d).

Overall, 121, 617, 667, and 674 genes were upregulated by acetic, propionic, butyric, and valeric acid treatment, representing a total of 1,031 unique genes (Figure 4c). Of these, 315 upregulated genes were common to valeric, butyric, and propionic acid exposure. GO analysis revealed that pathways related to fatty acid catabolism, fatty acid β-oxidation, and monocarboxylic acid catabolism were upregulated in this subset (Figure 4e). Moreover, analysis of the most differentially expressed genes (< or > 4 log_2_ change in expression, adjusted P value <0.001) highlighted an increase in genes implicated in modified β-oxidation, as evidenced by the upregulation of POX1-3 and PXP2 (Extended Figure 5). Since modified β-oxidation has been shown to metabolize toxic intermediates such as propionyl-CoA^39^, the *C. parapsilosis* transcriptome analysis supports the idea that SCFA exposure activated a detoxification mechanism.

**a**, Principal component analysis of global gene expression by *C. parapsilosis* cells treated with 0 mM (control, orange dots) or 25 mM acetic (light blue dots), propionic (green dots), butyric (red dots), or valeric acid (purple dots) for 90 min.

**b-c**, Venn diagrams depicting the number of downregulated (**b**) or upregulated genes (**c**) in *C. parapsilosis* cells treated with valeric acid (VA), butyric acid (BA), propionic acid (PA), or acetic acid (AA), compared to untreated control cells.

**d, e**, Gene Ontology (GO) analysis, based on differential gene expression in *C. parapsilosis* cells treated with VA, BA, or PA highlighted in (**b, c**), shows (**d**) the 6 most downregulated (black bars, −log_10_(adjusted *P* value) >40) and (**e**) the 5 most upregulated (white bars, −log_10_(adjusted *P* value) > 4) biological processes (BP) compared to control cells.

Statistical analysis: **b, c,** Differential gene expression analysis with adjusted *P* value was performed using DESeq2. Genes with log_2_(fold change) > 1 or < −1 with an adjusted *P* value < 0.05 were considered significantly differentially expressed. **d, e,** Gene Ontology analysis was performed using g:Profiler with g:SCS correction for multiple testing.

### Valeric acid disrupts *C. parapsilosis* intracellular pH homeostasis

To test the idea that valeric acid causes *C. parapsilosis* growth inhibition in a pH-dependent manner, we neutralized the growth medium supplemented with valeric acid (*pKa* = 4.84 at 25 °C) to a pH value of 7.0 with HCl. pH neutralization impaired the *Candida* growth-inhibitory effect of valeric acid. However, acidification of YPD broth to pH 5.5 (with HCl), the same pH value observed with addition of 6 mM valeric acid, did not inhibit *C. parapsilosis* or *C. metapsilosis* growth (Figure 5a-5b and Extended Figure 6a). These results indicate that valeric acid-mediated growth inhibition occurs at an acidic pH which favors the protonated, uncharged species of the monocarboxylic acid.

**Figure 5.**
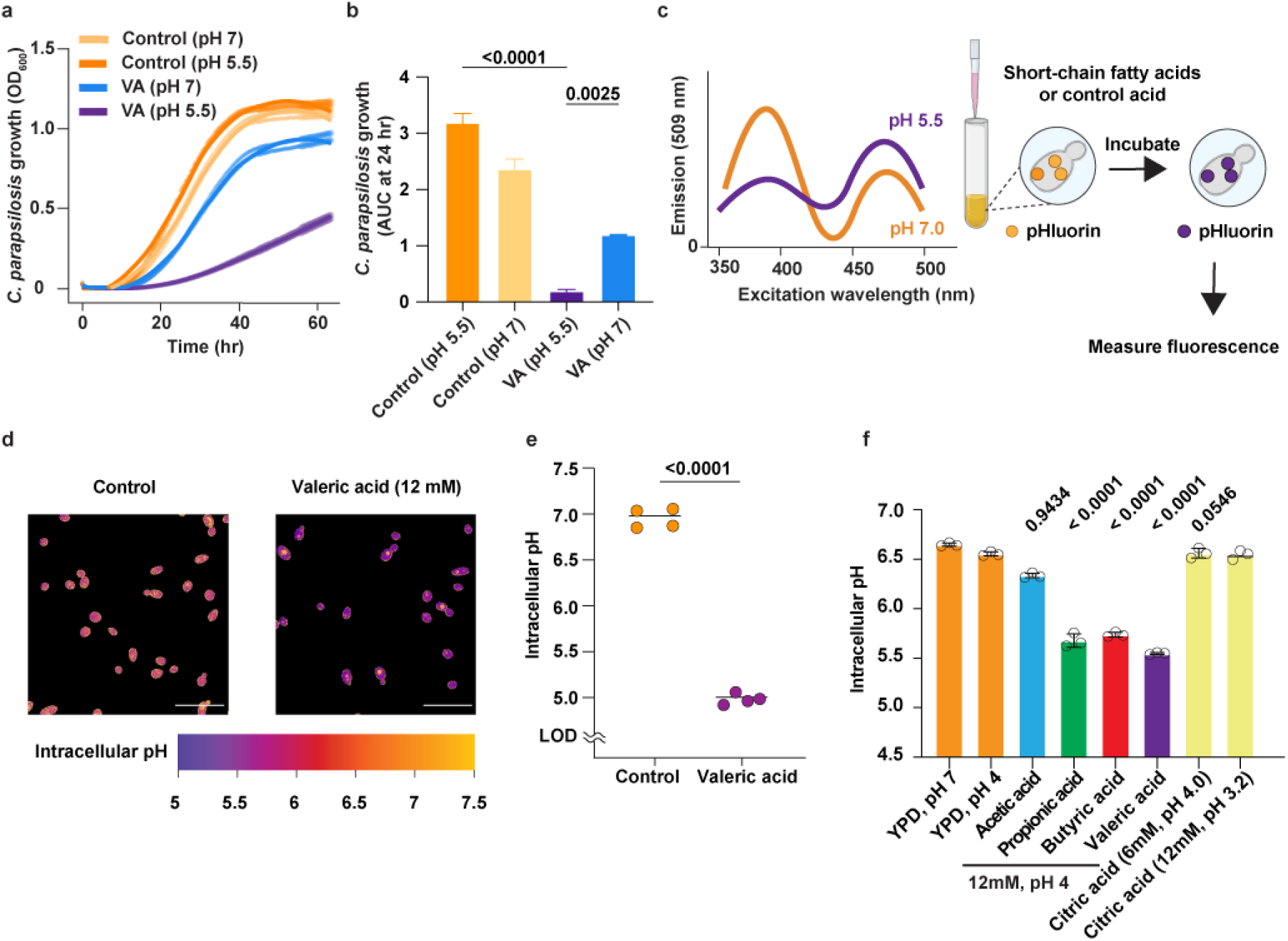
Valeric acid exposure and loss of *C. parapsilosis* pH homeostasis. **a, b,** *C. parapsilosis* 1004 (**a**) growth kinetics by optical density (600 nm) and (**b**) AUC values at 24 hr after culture in YPD broth with 0 mM (pH 5.5, orange symbols or pH 7, yellow symbols) or with 6 mM valeric acid (pH 5.5, purple or neutralized to pH 7, blue symbols). Data are representative of at least two independent experiments and growth curves were calculated from at least two replicates in an experiment. **c,** pHluorin emission spectrum at pH 5.5 (purple line) and pH 7 (orange line) and experimental scheme with pHluorin-expressing *C. parapsilosis* cells. **d-f**, pHluorin-expressing *C. parapsilosis* 65 was incubated in YPD broth with the indicated concentrations (0-12 mM, 0 mM control, orange) of (**d-f**) valeric acid (purple symbols), (**f**) acetic acid (light blue symbols), (**f**) propionic acid (green symbols), (**f**) butyric acid (red symbols) or (**f**) citric acid (yellow symbols) followed by (**d, e**) fluorescence microscopy emission quantification at 525 nm with 405 and 488 excitation wavelengths or (**f**) fluorescence plater reader emission quantitation at 509 nm with 395 and 475 nm excitation wavelengths. Representative images or data from two independent experiments shown and the intracellular pH is indicated using a standard curve of permeabilized pHluorin-expressing cells by monensin. (**d**), Scale bar, 20 µm. Statistical analysis: **b**, Adjusted *P* values versus control, calculated with 2-way ANOVA with Bonferroni’s multiple comparison test. **e**, Welch’s two-sided test: *P* < 0.0001 (n = 4). **f**, Data as mean values are shown with adjusted *P* values versus pH-adjusted control or indicated pairs (*P* value: 12 mM of indicated short-chain fatty acid vs control pH 4, or 6 mM of citric acid vs control pH 4) calculated with 2way ANOVA with Bonferroni’s multiple comparison test.

The concept of intracellular acidification as a mechanism of bacterial growth inhibition was discovered in bacterial pathobionts, specifically *Salmonella* and *Enterococcus*^40, 41^. To explore this idea in fungi, we generated *C. parapsilosis* strains that express a GFP-based ratiometric pH sensor, termed pHluorin (Figure 5c)^42, 43, 44, 45, 46, 47,48^. The pHluorin-expressing *C. parapsilosis* strains exhibited no growth defect compared to the parental strains and a standard pH reporter curve was generated using permeabilized cells (Extended Figure 6b-6c).

**Figure 6.**
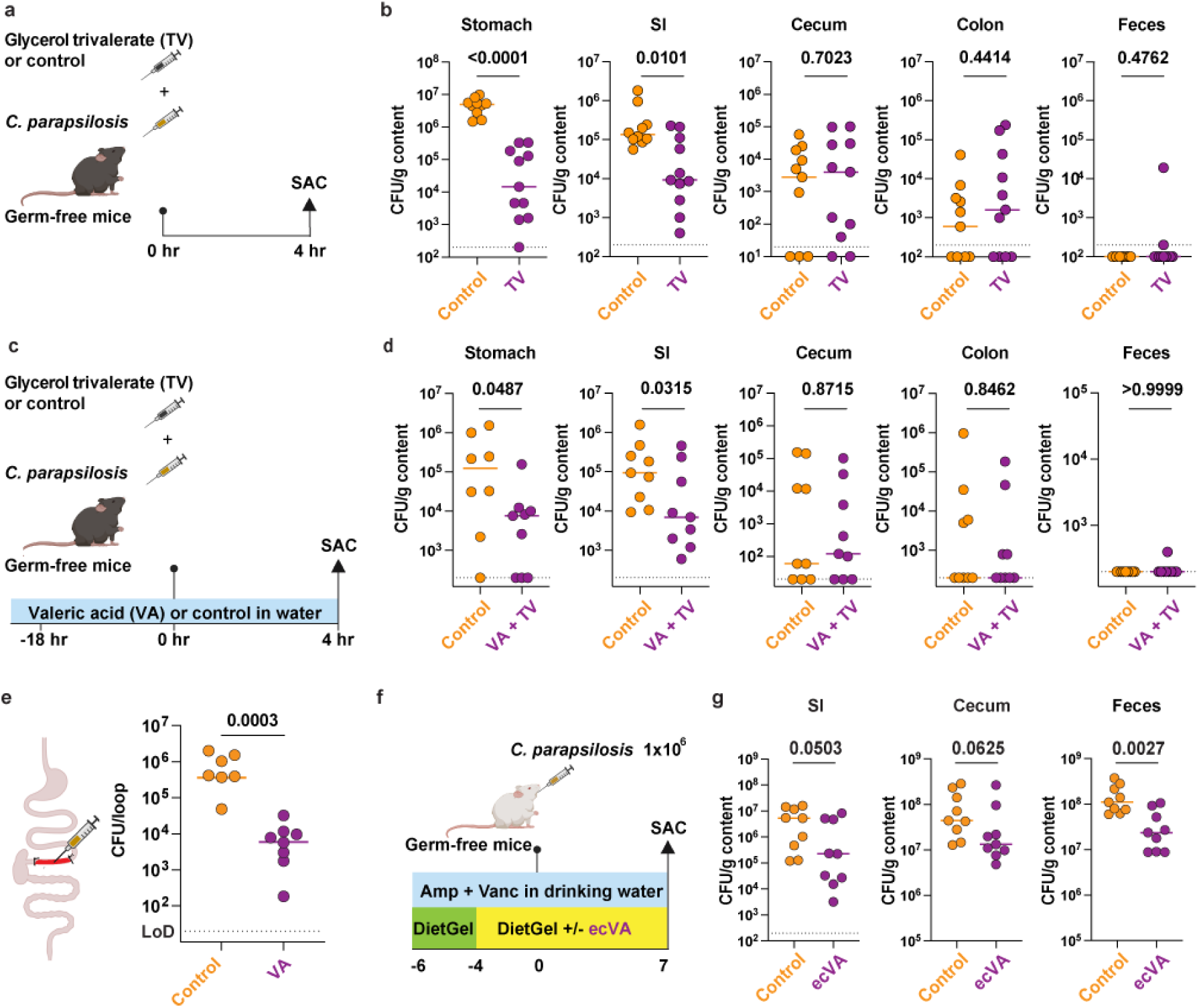
Valeric acid-associated inhibition of *C. parapsilosis* intestinal expansion. **a, c, e, f,** Experimental schemes to test the administration of (**a, c**) glycerol trivalerate (TV, gavage), (**c, e**) valeric acid (VA, **c,** drinking water *ad libitum* and **e**, injection into an ileal loop), and (**f**) microencapsulated valeric acid (ecVA) by addition to DietGel93 feed on *C. parapsilosis* intestinal colonization. **b, d,** *C. parapsilosis* CFU in stomach, small intestine, cecal, or colonic contents, or in the feces of germ-free C57B/6J mice that received 200 µl of 1M TV (purple dots) or glycerol (control, orange dots) and 1 x 10^6^ *C. parapsilosis* by gavage, with mice maintained on 0 mM (**b**, both groups; **d**, control group treated with glycerol) or 150 mM VA in the drinking water (**d**, TV group). **e,** Ileal loop *C. parapsilosis* CFU 3 hr after injection with 300 μl solution that contained 10^5^ *C. parapsilosis* cells and 0 (control) or 100 mM valeric acid in PBS, adjusted to the same pH. **g,** *C. parapsilosis* CFU in stomach, small intestine (SI), cecal, or colonic contents, or in the feces of germ-free, antibiotic-treated C57B/6J mice that received microencapsulated valeric acid or no capsules in feed starting on day −4 to day +7 after gavage with 10^6^ *C. parapsilosis* cells. (**b, d, g**) Results were pooled from two independent experiments or were (**e**) representative of 3 independent experiments, with 7-11 mice per group. Each dots represents one mouse. Statistical analysis: *P* values were calculated using two-tailed Mann–Whitney U test.

Treatment of pHluorin-expressing *C. parapsilosis* 65 with 12 mM valeric acid caused a decrease in mean intracellular pH from 6.96 to 5.00 (Figure 5d-5e), indicating a loss in intracellular pH homeostasis which is linked to a decrease in translational activity^48^. Using two independent pHluorin-expressing *C. parapsilosis* strains cultured in 6-12 mM 2-5 carbon chain SCFAs for 1 h, valeric acid caused the largest decrease in intracellular pH, with smaller changes observed with butyric and propionic acid (Figure 5f, Extended Figure 6d), and no change observed with acetic acid (Figure 5f, Extended Figure 6d). As a control, citric acid did not induce *C. parapsilosis* intracellular acidification even though it decreased the extracellular pH more than valeric acid and other tested SCFAs (Figure 5f, Extended Figure 6d). These results indicate that intracellular *C. parapsilosis* acidification is specific to certain SCFAs but not a common effect of exposure to other weak organic acids.

### Valeric acid treatment impacts intestinal *Candida* growth in mice

To test whether valeric acid could directly alter *C. parapsilosis* colonization *in vivo*, we infected germ-free C57B/6J mice with 10^6^ *C. parapsilosis* cells followed by treatment with glycerol trivalerate (TV), which consist of a glycerol backbone with three esterified valeric acid chains, and enumeration of CFU in the gastrointestinal tract 4 hr later (Figure 6a). Gastric and pancreatic lipases can hydrolyze TV to release free fatty acids^49, 50, 51, 52, 53, 54^. TV reduced *C. parapsilosis* colonization in the stomach and small intestinal contents but did not in the lower intestinal contents or feces, compared to treatment with a glycerol control (Figure 6b).

To deliver of valeric acid throughout the intestinal tract, we next administered 150 mM valeric acid in the drinking water overnight to germ-free C57BL6/J (Figure 6c) or Balb/c mice (Extended Figure 7a), followed by gavage of 1×10^6^ of *C. parapsilosis* and TV or control wvehicle with euthanasia 4 h later. Receipt of valeric acid and TV reduced *C. parapsilosis* colonization in the stomach and small intestines. However, there was no significant difference in *C. parapsilosis* CFU in lower intestinal segments, as previously observed with TV administration alone (Figure 6d, Extended Figure 7a-7b).

To determine whether the reduction of *C. parapsilosis* CFU in the stomach and small intestine was linked to local valeric acid concentrations, we measured valeric acid concentrations in the gastric, small intestinal, cecal, and large intestinal contents. We detected valeric acid in stomach and small intestinal contents but not in cecal and large intestinal contents (Extended Figure 7c-7d). These results indicate that valeric acid and TV supplementation increased valeric acid concentrations only in the stomach and small intestine, and that this localized increase correlated with the reduction in *C. parapsilosis* colonization (Figure 6c-6d, Extended Figure 7a-7d).

To test whether valeric acid could directly reduce *C. parapsilosis* levels in the lower intestinal tract, we developed an externalized ileum loop model in which *C. parapsilosis* (1 x 10^5^ cells) and valeric acid were co-administered. Direct delivery of valeric acid to the ileum inhibited the growth and expansion of *C. parapsilosis* CFU by ∼2log_10_ in the ileal lumen 3 h after murine infection (Figure 6e).

To release valeric acid along the entire intestinal tract, we encapsulated and coated valeric acid through a spray cooling-based microencapsulation and mixed the resulting capsules into DietGel93M (at a final calculated valeric acid concentration of 88.6 mM) for oral administration *ad libitum* in germ-free mice (See Materials and Methods)^55^. Germ-free mice were started on vancomycin and ampicillin on day −6 to prevent any bacterial growth. Mice received valeric acid capsules or no capsules on day - 4 and were administered 1 x 10^6^ *C. parapsilosis* cells on day 0 (Figure 6f). Receipt of microencapsulated valeric acid in the diet reduced *C. parapsilosis* CFU in the lower intestinal contents and feces on day 7 (Figure 6g). In parallel, we measured valeric acid concentrations in intestinal contents on day +7 and detected valeric acid in the feces of most treated mice, but not in untreated mice (Extended Figure 6e). Collectively, these results indicate that valeric acid can inhibit intestinal *Candida* growth at sites where the compound can be delivered and detected.

## Discussion

In this study, we employed a machine learning approach to identify a bacterial-derived metabolite, valeric acid, that mediates trans-kingdom ecology, inhibiting *C. parapsilosis* growth. This was validated experimentally through analysis of human allo-HCT fecal samples, *in vitro* growth assays, and *in vivo C. parapsilosis* colonization studies in mice. Valeric acid exerted growth-inhibitory effects across a broad range of *Candida* species, including *C. albicans* and *C. auris*, and was more potent in growth inhibition than propionic and butyric acid. Valeric acid resulted in loss of intracellular pH homeostasis in *C. parapsilosis* and *C. albicans* (Mishra et al, companion study) and this disruption was linked to a reduction in protein synthetic ability and fungal cell growth. Loss of intracellular pH homeostasis represents can be exploited as an antifungal strategy^48^. For example, antifungal terpenoid phenols cause a burst of Ca^2+^ transients, disrupting intracellular pH homeostasis, followed by repression of genes mediating ribosome biogenesis and RNA metabolism^56^.

SCFA-mediated loss of intracellular pH homeostasis is conserved across microbial kingdoms and acts on bacterial pathobionts as well, exemplified by studies on *Salmonella*^57^ and *Enterobacteriaceae*^40^. This work and the companion study by Mishra *et al*. extends this concept to pathogenic eukaryotes, establishing it as a key mechanism of trans-kingdom ecology of the intestinal microbiota. In a *C. albicans* study, exposure to 18 mM butyric acid, 52 mM propionic acid, and 75 mM acetic acid led to downregulation of genes involved in RNA synthesis and ribosome biogenesis as well as a reduction in total RNA extracted per unit optical density of *C. albicans* cultures^30^. In other studies, SCFA exposure impacted *C. albicans* gene metabolism by regulating histone acylation and chromatin structures, which activate or repress transcription and thus affect cell wall structure and yeast-to-hypha morphogenesis, biofilm formation, and antifungal drug susceptibility^28, 34, 58^.

The mechanism by which SCFAs enter fungal cells remains poorly understood. Monocarboxylate transporters, such as Jen1, or acetate transporter families, such as Ady2, have been characterized in *S. cerevisiae*, and orthologs have been identified in *Candida* species^59, 60, 61^. Although no specific transporters have been identified as SCFA transporters in *Candida,* it is possible that variations in transporter specificity or expression in different *Candida* species may explain the susceptility of different *Candida* species or isolates to growth-inhibitory effects of SCFAs. In addition, protonated SCFAs may passively diffuse into fungal cells^62^.

The growth-inhibitory effects of SCFAs of varying length may also be related to differences in intracellular metabolism and the potential toxicity of metabolic intermediates.The accumulation of propionyl-CoA, 3-hydroxypropionate, and 3-hydroxypropionyl-CoA exerted toxicity in both host and fungal cells^63, 64, 65^. Activation of the modified beta-oxidation pathway may reflect a protective mechanism for detoxifying these intermediates by converting them into acetate and acetyl-CoA. This may partially explain the lack of growth-inhibitory effects of acetic acid at the concentration range tested. Valeric acid may also generate toxic intermediates, including propionyl-CoA, through canonical beta-oxidation^39^, contributing to its inhibitory effect. In contrast, butyric acid undergoes canonical beta-oxidation to form acetyl-CoA, yet still effectively inhibits *Candida* growth, suggesting that propionyl-CoA accumulation alone may not fully explain SCFA-mediated growth inhibition. Further research is warranted to elucidate the intracellular metabolism and degradation of internalized short chain fatty acids.

In allo-HCT patients, intestinal *Candida* expansion is linked to poor clinical outcomes, in part due to heightened risk of bloodstream translocation^14, 15^. In allo-HCT patient fecal samples, *C. papapsilosis* expansion occurred only when SCFA abundances were low. *C. parapsilosis* expansion did not occur in all samples with low SCFA abundances, due to potential loss of *C. parapsilosis* from the intestinal microbiota prior to the reduction in SCFA abundance or due to the presence of alternative mechanisms of colonization resistance, e.g., host-derived cathelicidins, hypoxic tissue conditions, the presence of bacterial competitors, or nutrient limitation^18, 24, 66^. Receipt of medications that regulate pH along the gastrointestinal tract likely influences the relationship of SCFA abundance and *Candida* colonization as well^67^. Fecal metabolite analysis in this study was limited in scope and fecal samples with low SCFA concentrations may have contained other compounds that prevented *C. parapsilosis* growth. For example, we observed that indole abundance inversely correlated with *C. parapsilosis* growth. Larger untargeted metabolomic studies will be required to identify and test additional host- and bacterial-derived metabolites implicated in *Candida* growth inhibition.

Beyond lowering *C. parapsilosis* colonization, administration of glycerol trivalerate to mice can inhibit the growth of the bacterial pathogen *Clostridiodes difficile*^54^. In a study of antibiotic-induced microbiota states that promote *C. difficile* colonization, administration of cefoperazone reduced intestinal valeric acid levels by 66-fold^68^. Collectively, these studies link changes in valeric acid abundance to *C. difficile* infectious susceptibility in mice. Intestinal valeric acid abundance has been linked to host immune activation by regulation of mammalian target of rapamycin activity in B cells and CD4^+^ T cells, with effects on IL-17 and IL-10 release by effector T cell populations^69, 70^. Butyric acid is a well-known inducer of colonic colonic T regulatory cell differentiation^71^. Thus, it is possible that valeric acid (and other SCFAs) could exert effects on bacterial and *Candida* intestinal colonization through indirect, immune-mediated mechanisms or through effects on commensal bacteria (Mishra et al., companion study).

Some commensal bacterial species, including *Megasphaera massiliensis, Anaerostipes hadrus,* and *Coprococcus comes*, produce valeric acid, and its physiological intestinal concentration is typically lower than that of other SCFAs^69^. Since valeric acid is a potent SCFA against *Candida* species, and its physiological concentration is in the low mM range at baseline, it is possible that strategies to promote a modest increase in valeric acid concentration may be effective in limiting *Candida* growth. Patients with leukemia and solid organ and bone marrow transplant recipients are highly vulnerable to systemic candidiasis and commonly receive prophylactic antibiotics to prevent life-threatening bacterial infections. Thus, strategies that depend on live commensal bacteria (i.e., fecal microbiota transplantation or delivery of rational bacterial consortia) to reverse microbial dysbiosis are limited by the ongoing need for antibacterial antibiotics during high-risk periods. The strategy of using metabolites to mitigate pathobiont expansion is attractive in these clinical scenarios. A major barrier to clinical success remains the development of strategies to deliver metabolites to intestinal sites of pathobiont expansion. Our findings support the idea that interventions to increase intestinal valeric acid levels may be beneficial for immunocompromised patients with dysbiosis, such as allo-HCT patients, in whom *Candida* expansion and bloodstream infection originating from intestinal translocation impacts clinical outcomes.

## Acknowledgments

We thank Dr. Julia Köhler (Boston’s Children’s Hospital, Harvard University, USA) for providing the pHluorin construct, the MSK Imaging core facility for microscopy experiments, the Integrated Genomics Operation Core for RNA-seq experiments. Graphical schemes were created with BioRender.com. Portions of this text were edited with the assistance of ChatGPT (OpenAI, 2025). The resulting content was reviewed and further edited by the authors. These studies were supported by a Geoffrey Beene Foundation Award (TMH), a Burroughs Wellcome Fund Investigator in the Pathogenesis of Infectious Diseases Award (TMH) and by NIH grants P01 AI179406 (TMH and JBX), R37 AI093808 (TMH), K99 AI175599 (CL), P30 CA008748 (to MSKCC), an unrestricted award from Tito’s Love, a Takeda Science Foundation Fellowship grant (KYM), and JSPS KAKENHI Grant JP202470035 (KYM).

## Author Contributions

The study was conceived by T.M.H. and K.Y-M. *In vitro* experiments and *in vivo* experiments were performed by K. Y-M., A. B and C.N.S. *C. parapsilosis* pHluorin strains were generated by T. N and A.G. Transcriptome analysis was performed by K.Y-M, K.B., and G.B. Imaging analysis was performed by E.C. and K.Y-M. SHAP modeling analysis was performed by C.L. and J.B.X. GC-MS analysis was performed by R.R., J.C., A.S, and E.G.P. Microencapsulated valerate was generated by P.M. Data analysis and visualization were performed by K.Y-M. and C.L. Data interpretation was performed by K.Y-M. and T.M.H. The manuscript was written by K.Y-M. and T.M.H. All authors have read and approved the manuscript.

## Resource availability

### Lead Contact

Further information and requests for resources and reagents should be directed to the lead contact, Tobias Hohl.

### Materials Availability

Requests for materials should be addressed to T.M.H and should be covered by a material transfer agreement.

### Data and Code Availability

The *C. parapsilosis* RNA-Seq dataset is being deposited in the Sequence Read Archive under BioProject PRJNA1294161 (reviewer link at URL https://dataview.ncbi.nlm.nih.gov/object/PRJNA1294161?reviewer=fiaeishv6qhuim2sk35rrham0t). Fungal ITS1 and bacterial 16S amplicon sequences previously were deposited under BioProject PRJNA746305 and re-analyzed for the relevant patient fecal samples (see Supplementary Table 4)^14^. All data needed to evaluate the conclusions are present in the paper or in the Supplementary Materials. Any additional information required to re-analyze the data reported in this paper is available from the lead contact upon request. All code for this study is at https://github.com/liaochen1988/Cparaplosis_gut_colonization.

## Methods

### Bacteria and *Candida* strains

Commensal bacterial strains isolated from healthy human donor feces were stored in the MSKCC BioBank or the Symbiotic Strain Bank at Duchossois Family Institute at the University of Chicago. *C. parapsilosis* complex strains are MSK clinical isolates, with *C. parapsilosis* strains 1004 and 1666 and *C. metapsilosis* strains 73 and 414 from fecal samples and *C. parapsilosis* strains 478, 65, and 804, *C. orthopsilosis* strains 479 and 616, and *C. metapsilosis* strain 798 from blood culture specimens^72^. *C. auris* strain 1193 was isolated from human nares. *C. albicans* strains SC5314^73^, 529L^74^, CHN1^75^, *C. glabrata* (ATCC 2001), and *C. tropicalis* (ATCC750) were from the American Tissue Culture Collection (ATCC). All fungal and bacterial strains isolated at MSKCC or at the University of Chicago were classified based on rDNA (16S rDNA for bacteria or ITS1 for fungi) or on whole genome sequencing.

### *In vitro* screen of *C. parapsilosis* growth in bacterial spent supernatants

Individual bacterial strains were shaken in 96 well plates at 37°C in brain heart infusion liquid medium (BHI; BD cat no. 237500) supplemented with 0.5 g/L cysteine, 1 mg/L menadione, and 10 mg/L hemin) under anaerobic conditions to yield stationary phase cultures and spent supernatants were collected by centrifugation. An overnight culture of *C. parapsilosis* strain 1004 was grown in yeast extract peptone dextrose (YPD) broth (BD cat no. 242820) overnight, washed twice in PBS, and diluted to add 400 CFU to 100 μl fresh BHI broth mixed with 100 µl of spent bacterial supernatant or with 100 μl PBS (control) in a 96-well plate. The cultures were incubated in a SpectraMax i3 Microplate Reader (Molecular Devices, CA, USA) at 30°C with shaking under aerobic conditions. Area under the curves (AUC) for 24 h of growth was calculated using Growthcurver in R. To assess the effect of specific metabolites on *Candida* growth, indicated *Candida* strains were inoculated into 96-well plates at 400 CFU/well in YPD broth supplemented with the indicated metabolite concentrations, and growth curves were generated in a SPARK microplate reader (Tecan, Switzerland).

### Fecal sample collection in MSK allo-HCT patients

MSK adult allo-HCT patients were invited to participate in an IRB-approved longitudinal collection of fecal samples as described^14, 15^. Per institutional guidelines, allo-HCT patients received micafungin (100 mg IV every 24 hr) and levofloxacin (500 mg po every 24 hr) for prophylaxis, starting at the beginning of pre-transplant conditioning chemotherapy. For allo-HCT patients who received total body irradiation as part of the conditioning regimen, vancomycin (1 gm IV every 12 hr) was administered from day −2 to day + 7. Micafungin was switched to a mold-active azole drug (i.e, voriconazole or posaconazole) on day +7 for unmodified myeloablative, cord blood, haploidentical, and T cell-depleted allo-HCT. In this study, we included 145 fecal samples from 22 patients with at least one culture-positive *C. parapsilosis* sample per patient and one culture-negative sample per patient. 56 samples from these 22 patients were excluded from analysis because of (1) concurrent receipt of triazole therapy or (2) sample culture positivity for fungi other than *C. parapsilosis complex* species.

### Metabolomic analyses

Targeted metabolomics was performed to determine the abundance of 38 metabolites in spent bacterial supernatants at University of Chicago or the abundance of 43 metabolites including short-chain fatty acids in human and mouse fecal samples at MSKCC.

#### Supernatant sample extraction and derivatization

Supernatants were extracted and derivatized as described^76, 77^. Briefly, supernatants were extracted using 4 volumes of 100% methanol spiked with internal standards, centrifuged at 20,000 x *g* for 20 min at 4°C.

#### Human and mouse sample extraction

Human fecal samples and the murine small intestine, cecum, colon, and fecal samples were weighed and transferred into 2 mL microtubes pre-filled with 1.4 mm ceramic beads (Omni International). Human and murine fecal samples were resuspended to a final concentration of 100 mg/mL and 50 mg/mL in 9 volumes of 80:20 methanol:water (Optima LS/MS grade #A456, Fisher Scientific, MA) that contained acetate-d3, propionate-d5, butyrate-d7, and valerate-d9 as internal standards (Cambridge Isotope Laboratories). Homogenization was performed using a Bead Ruptor (Omni International) at a speed of 6 m/s for 3 minutes at 4°C. After homogenization, the samples were centrifuged at 20,000 x *g* for 20 minutes at 4°C to collect the supernatant.

#### Sample derivatization and quantification by GC-MS

Extracted supernatant, human and mouse samples were derivatized for metabolomic analysis similar to methods as described^76^ with modifications^77^. Briefly, 1 volume of extract was added to 1 volume of 100 mM borate buffer (pH 10), 4 volumes of 100 mM pentafluorobenzyl bromide (PFBBr, Thermo Scientific) diluted in acetonitrile (Fisher), and 4 volumes of n-hexane or cyclohexane (Acros Organics) and quantified by gas-chromatography-mass spectrometry (Agilent 7890A GC and Agilent 5975C MS detector) operating in negative chemical ionization (nCI) mode, using methane as the reagent gas. For supernatant analysis, semi-quantitative values were calculated from the raw peak area of the compound normalized to the median raw peak area of two baseline media controls. For human and mouse SCFA analysis, the raw peak areas of acetate (*m/z* 59) and propionate (*m/z* 73) were normalized to acetate-d3 (*m/z* 62) and propionate-d5 (*m/z* 78), respectively; butyrate and isobutyrate (*m/z* 87) were normalized to butyrate-d7 (*m/z* 94); 2-methylbutyrate, valerate, and isovalerate (*m/z* 101) were normalized to valerate-d9 (*m/z* 110). Data was analyzed using the MassHunter Quantitative Analysis software (Version 10.1, Agilent Technologies) and confirmed by comparison to authentic standards.

### Random Forest model and SHAP analysis

A Random Forest (RF) model of 1,000 trees was constructed to predict *C. parapsilosis* growth based on metabolite concentrations in bacterial supernatants. Two metabolites were excluded from the analysis due to insufficient data points. Metabolite profiling was available for 229 out of 347 isolates used in growth analysis, and these 229 isolates with data on 36 metabolites were used for modeling and SHAP analysis. Both metabolite concentrations and growth curve AUCs were log2-transformed. The RF model hyperparameters were fine-tuned to maximize the R2 score through a grid search in a 5-fold cross-validation framework. These hyperparameters include the number of features considered for the optimal split (max_features = ‘sqrt’, ‘log2’, 0.25, 0.50, 0.75, 1.00), the maximum depth of each tree (max_depth = 2, 4, 8), the minimum number of samples required to split an internal node (min_samples_split = 2, 4, 8), and the minimum number of samples required at a leaf node (min_samples_leaf = 1, 2, 4). The RF model was implemented using the RandomForestRegressor function from Python’s scikit-learn package (version 1.2.2). SHAP (SHapley Additive exPlanations) analysis was performed using the TreeExplainer function from the SHap package (version 0.42.0).

### pHluorin-expressing *Candida parapsilosis* strains, pH calculation, and visualization

The pHluorin-expressing construct was inserted into the CpNEUT5L locus of *C. parapsilosis* strains 65 and 804 by CRISPR-Cas9^42, 44^. The pHluorin ORF was amplified with primers pHluorin_ClaI_F (5’-ttt tat cga tat gtc aaa agg tga aga att att tac tgg-3’) and pHluorin_NheI_R (5’-ttt ttt tgc tag ctt att tat ata att cat cca tac cat gag taa tac c-3’). The amplicon and pNRVL-N5L-CFP were digested with *Cla*I and *Nhe*I and ligated to yield pNRVL-N5L-pHluorin (Accession number: PQ261162). This plasmid was used as a template to amplify the donor DNA with the primer pair FP_CpN5L_dDNA_F (5’-aac ctc atc tca agg cgc-3’) and FP_CpN5L_dDNA_R (5’-aca caa aaa tac atg att gcg tc-3’). CRISPR/Cas9 engineering and *C. parapsilosis* transformation of parental strains 65 and 804 was performed as described^47^ to generate pHluorin-expressing strains (genotype: *cpneut5l::PCaTDH3-pHLuorin-TScURA3/cpneut5l::PCaTDH3-pHLuorin-TScURA3*). The pHluorin-inserted Cp65 and 804 strains were cultured and inoculated into the media with SCFAs or calibration buffers and incubated for 1 h^43^. Fluorescence intensities were measured using a SPARK microplate reader (Tecan, Switzerland) at 509 nm emission with the excitation at 395 and 475 nm. Imaging was performed on a TCS SP5 confocal microscope (Leica, Germany) with 63x/1.4NA objective lens and fluorescence intensities were measured at 525 nm emission with the excitation at 405 and 488 nm. The image intensity was measured in FIJI (NIH, USA). Intracellular pH was determined based on fluorescence intensity using Graphpad Prism v9.0 and Fiji, based on the standard curve generated by monensin-permeabilizing cells.

### *Candida* RNA-seq analysis

*C. parapsilosis* strain 1004 was cultured at 30 °C overnight, diluted 1:100, and grown to an OD_600_ of 0.3-0.4 in YPD broth. After a PBS wash, 5 x 10^7^ cells were inoculated in 3 ml YPD with 0 or 25 mM SCFA and cultured for 1.5 h at 30°C with shaking, washed in PBS, stored in RNA later (Invitrogen, MA) at −80°C, thawed, and disrupted using RNA pink beads lysis kits (Next Advance, NY). RNA was isolated using TRIzol (Invitrogen, MA), cleaned with RNeasy (Qiagen, Netherland), quantified using RiboGreen (Thermo Scientific, MA), and analyzed for quality on an BioAnalyzer (Agilent, CA). 500 ng of total RNA with RIN values of 8.2-9.5 underwent polyA selection and used for TruSeq library preparation (TruSeq Stranded mRNA LT Kit, RS-122-2102, Illumina), with 8 PCR cycles. Barcoded samples were run on a NovaSeq 6000 in a PE100 run, using the NovaSeq 6000 S4 Reagent Kit (200 Cycles). Each sample generated an average of 52 million paired reads.

The raw data were analyzed using the Transcriptome pipeline (Basepair). Only genes with logFC >1 and p-value <0.05 in at least one comparison were included. Annotations for *C. albicans* and *S. cerevisiae* orthologs are taken from the *Candida* Gene Order Browser^78^. Orthologies were assigned on a single gene basis. For large gene families, only one *C. parapsilosis* gene was assigned as an ortholog. Gene annotation and clustering were performed using g:Profiler.

### Mice and treatment with valeric acid formulations

Specific pathogen-free (SPF) C57BL6/J male and female mice were used in experiments at 8-12 weeks of age. Germ-free C57BL6/J or Balb/c mice were bred and maintained in sterile isolators in the gnotobiotic facility at the Weill Cornell Medicine-MSKCC vivarium. All experiments complied with federal and institutional guidelines and were approved by the MSK Institutional Animal Care and Use Committee (Protocol #13-07-008).

Germ-free C57BL6/J or Balb/c mice were administered 150 mM valeric acid or control drinking water, both adjusted to pH 5, starting 18 hr prior to infection with *C. parapsilosis* strain 478 (inoculum: 10^6^ cells) and receipt of 200 μL of 1 M glycerol trivalerate or 1M glycerol in PBS, adjusted to pH 4.5, by oral gavage. Mice were euthanized 4 hr later and intestinal contents were harvested for quantitation of *Candida* CFU and valeric acid levels.

For the ileal loop model, SPF C57B6/J mice (Jackson Laboratories, strain 00664) were treated with ampicillin (1g/L) in the drinking water starting on day −3. On the day of the experiment, mice were anesthetized with ketamine and the ileum was externalized. The ileal loop was injected with 300 μl solution that contained 1×10^5^ *C. parapsilosis* strain 478 cells as well as 0 or 100 mM valeric acid in PBS adjusted to the same pH, mixed immediately prior to the injection. Mice were kept under anesthesia and on a heating pad for 3 hr prior to euthanasia for CFU enumeration from ileal loop contents.

Microencapsulated valeric acid was generated by SILA S.p.A. (Noale, Italy), according to the patent EP2352386B1 as follows^55^. Briefly, sodium valerate was generated by mixing valeric acid with an aqueous solution of 50% NaOH until the theoretical equivalence point was reached, ensuring complete conversion to sodium valerate while monitoring pH. The mixture was heated and dried at 120°C overnight, then ground into a fine powder. 41% (w/w) of sodium valerate was slowly added to a melted mixture of vegetable hydrogenated fatty acids and an emulsifier until a homogeneous dispersion was obtained, and the molten mixture was then cooled to room temperature to solidify. The solid product was ground with a low-shear mixer and sieved between 500 and 800 µm. The obtained titer was calculated to be 40.50%.

DietGel93M (ClearH2O, USA) was transferred to autoclaved feeders and mixed with or without valerate capsules and fed *ad libitum* to germ-free mice. To avoid any possible bacterial contamination, germ-free mice were maintained on ampicillin and vancomycin during the experiment and confirmed to be germ-free prior to *C. parapsilosis* administration.

Fecal, intestinal, and ileal loop contents were plated on SAB agar (BD Difco Sabouraud dextrose agar, BD 210930) plates supplemented with or without 10 μg/ml of vancomycin (Hospira, NDC 0409-6510-01) and 100 μg/ml of gentamicin (Gemini, 400108) as previously described^79^.

### Data analyses and plotting

GraphPad Prism (v9 or v10, GraphPad Software Inc.) or R (version 4.1.0, R Development Core Team) were used to generate data plots and perform statistical tests on datasets.

## Supplementary Information

**Supplementary Table 1. Bacterial supernatant metabolite profiles** (separate file due to size). Abundance of 36 target metabolites in spent supernatants of the 229 commensal bacteria from the library, compared to the BHI growth medium.

**Supplementary Table 2.** Clinical profiles of patients who provided fecal samples for analysis. Clinical background data of allo-HCT patients with fecal samples that were included in the study (n = 22).

|  | Overall |
| --- | --- |
| Number | 22 |
| Age (mean (SD)) | 57.33 (10.47) |
| Sex = F (%) | 12 (54.5) |
| Disease (%) |  |
| Hodgkin's Disease | 1 (4.5) |
| Leukemia | 10 (45.5) |
| Leukemia (Second Primary) | 1 (4.5) |
| Multiple Myeloma | 2 (9.1) |
| Myelodysplastic Syndrome | 2 (9.1) |
| Myeloproliferative Disorder | 2 (9.1) |
| Myeloproliferative Disorder (Second Primary) | 2 (9.1) |
| Non-Hodgkin's Lymphoma | 2 (9.1) |
| Source (%) |  |
| Allo BM Unmodified | 3 (13.6) |
| Allo PBSC CD34+ (CliniMacs/Miltenyi) | 13 (59.1) |
| Allo PBSC Unmodified Only | 6 (27.3) |

**Supplementary Table 3. Transcriptome analysis of SCFA-treated *C. parapsilosis* cells** (separate file due to size). List of downregulated or upregulated genes of *C.parapsilosis* cells treated with acetic, propionic, butyric or valeric acid compared to the untreated cells.

**Supplementary Table 4. 16S and ITS Sequences relevant for** Figure 2a. The data were previously deposited under BioProject PRJNA746305.

## Extended Data Figures

**Extended Figure 1.**
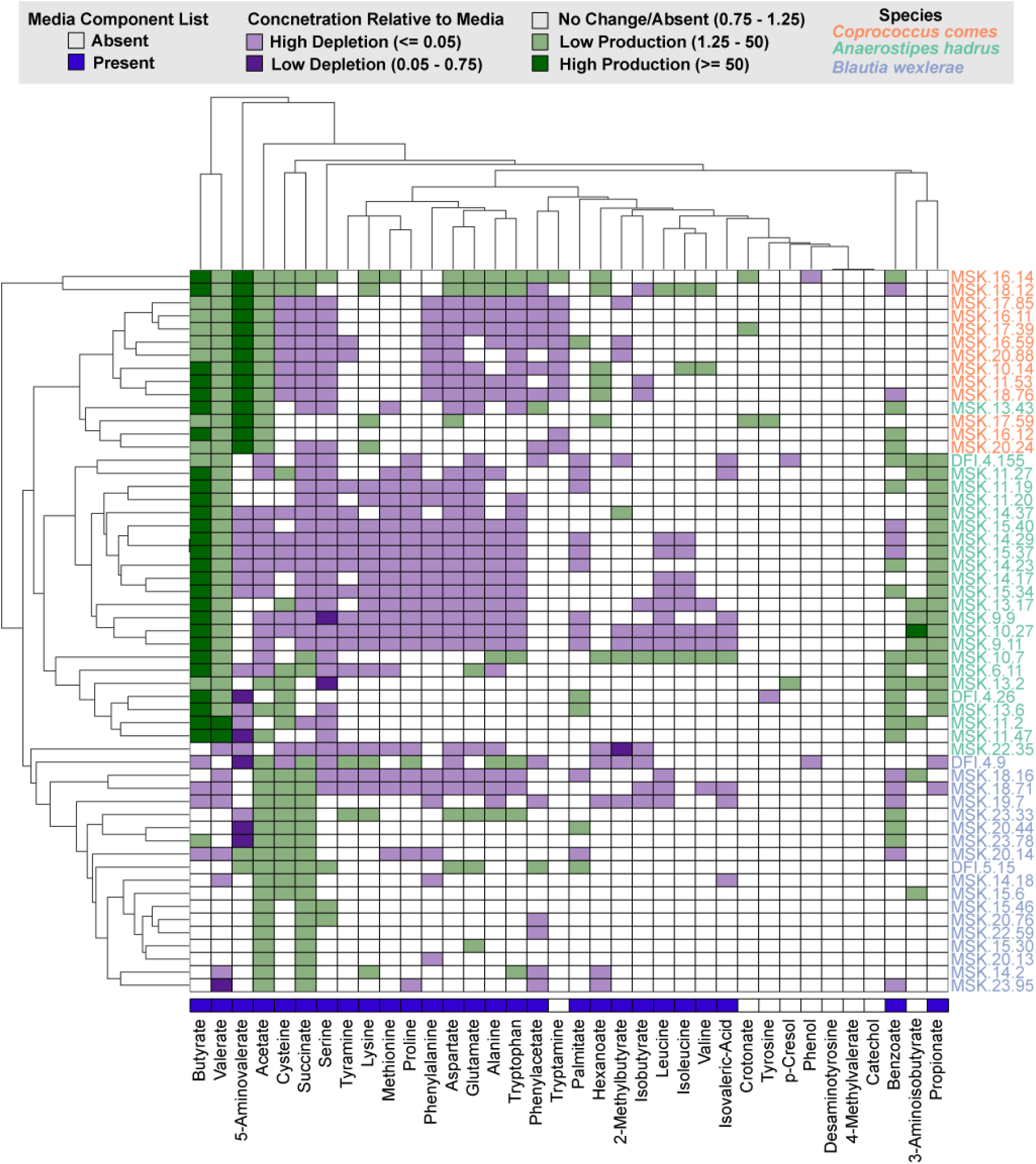
Heatmap of bacterial supernatant metabolite profiles, related to Figure 1. Concentrations of 36 metabolites in the spent supernatants of *Coprococcus comes* (orange), *Anaerostipes hadrus* (green), and *Blautia wexlerae* (blue) isolates were measured and shown relative to control BHI growth medium. Metabolite production is indicated in green and depletion in purple. Metabolites present in BHI growth medium are highlighted in blue.

**Extended Figure 2.**
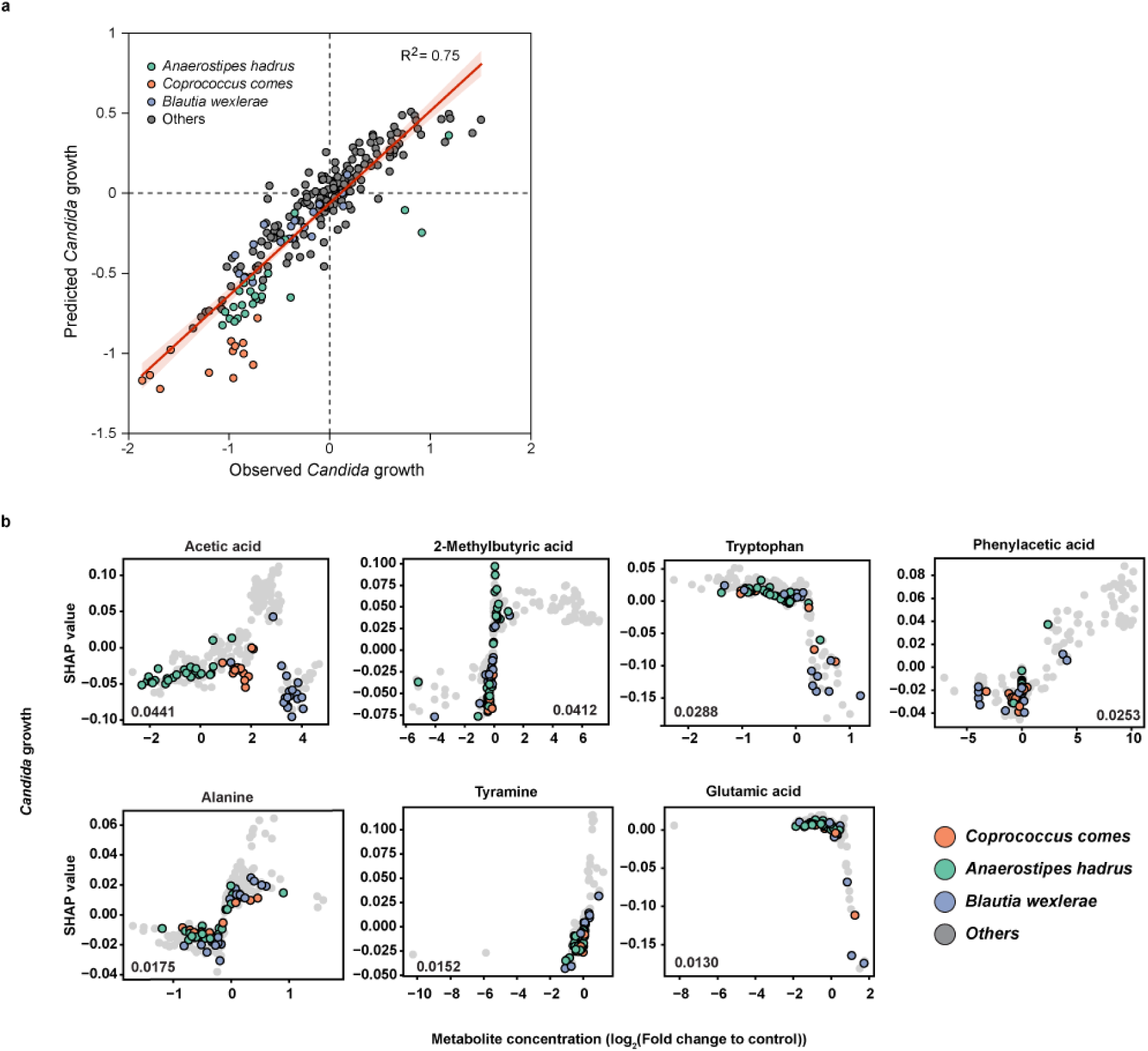
Prediction of bacterial metabolites that inhibit *C. parapsilosis* growth, related to Figure 1. **a,** Goodness of fit for the Random Forest model in reproducing the observed *Candida* growth, measured as area under the curve (AUC) relative to the negative control. **b,** SHAP analysis of *Candida* growth AUCs (y axis) in relation to metabolite concentrations (x-axis), including acetic acid, 2-methylbutyric acid, tryptophan, phenylacetic acid, alanine, tyramine, and glutamic acid. Each dot represents one bacterial isolate. *Coprococcus comes*, *Anaerostipes hadrus*, and *Blautia wexlerae* isolates are highlighted in orange, green, or blue, respectively. Figure 1c and Extended Figure 2b display the top ten metabolites, ranked by mean absolute SHAP values. SHAP values for all 36 metabolites, including the top ten.

**Extended Figure 3.**
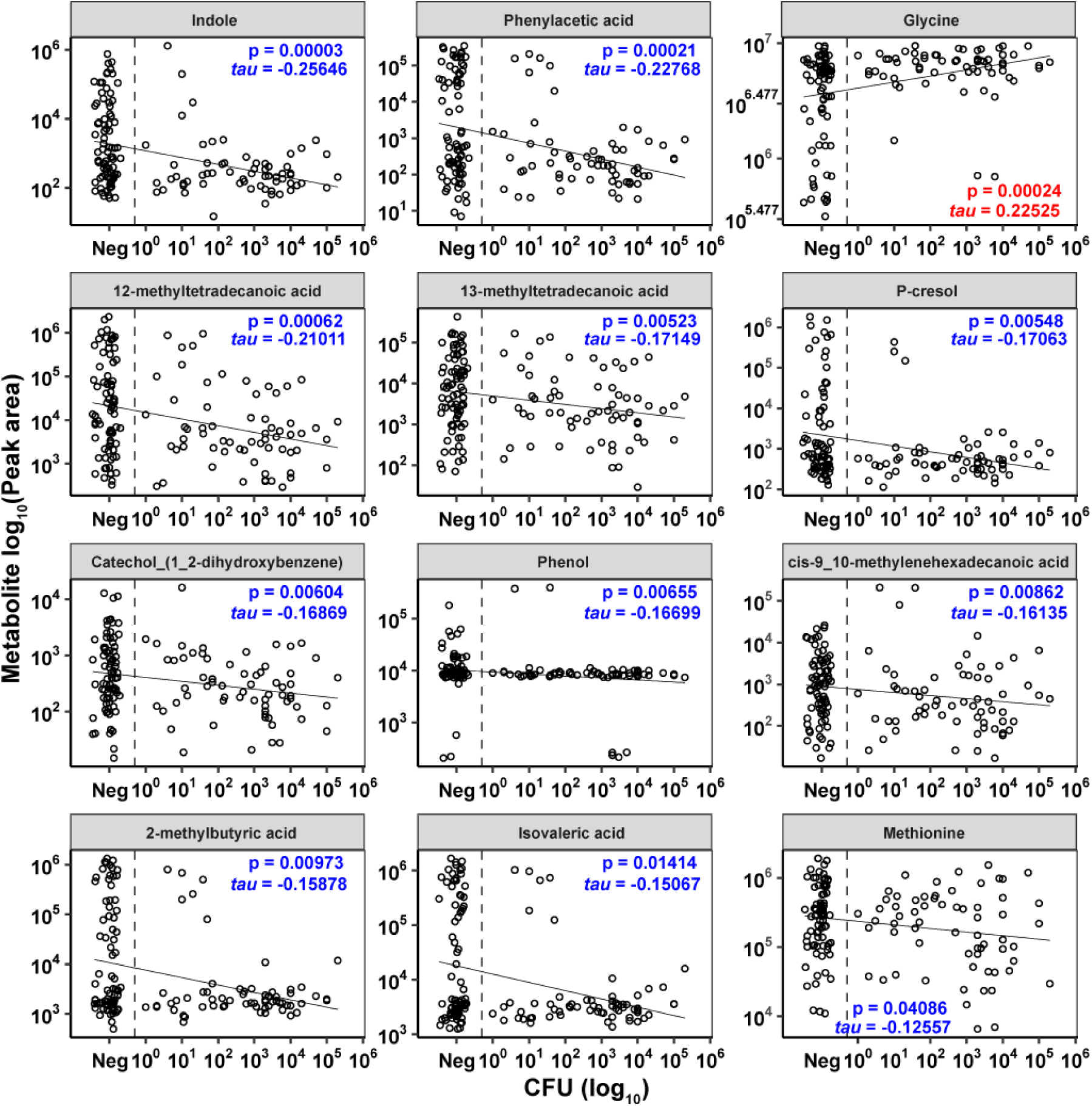
Allo-HCT patient fecal metabolite peak areas and *C. parapsilosis* growth, related to Figure 2. The plots indicate *C. parapsilosis* CFU and the GC/MS peak area of indicated metabolites in each sample (n = 145 fecal samples from 22 allo-HCT patients). Statistical analysis: Kendall’s *tau* and two-tailed *P* values are shown in blue (negative correlation) or in red (positive correlation). The plots with *P* <0.05 are shown.

**Extended Figure 4.**
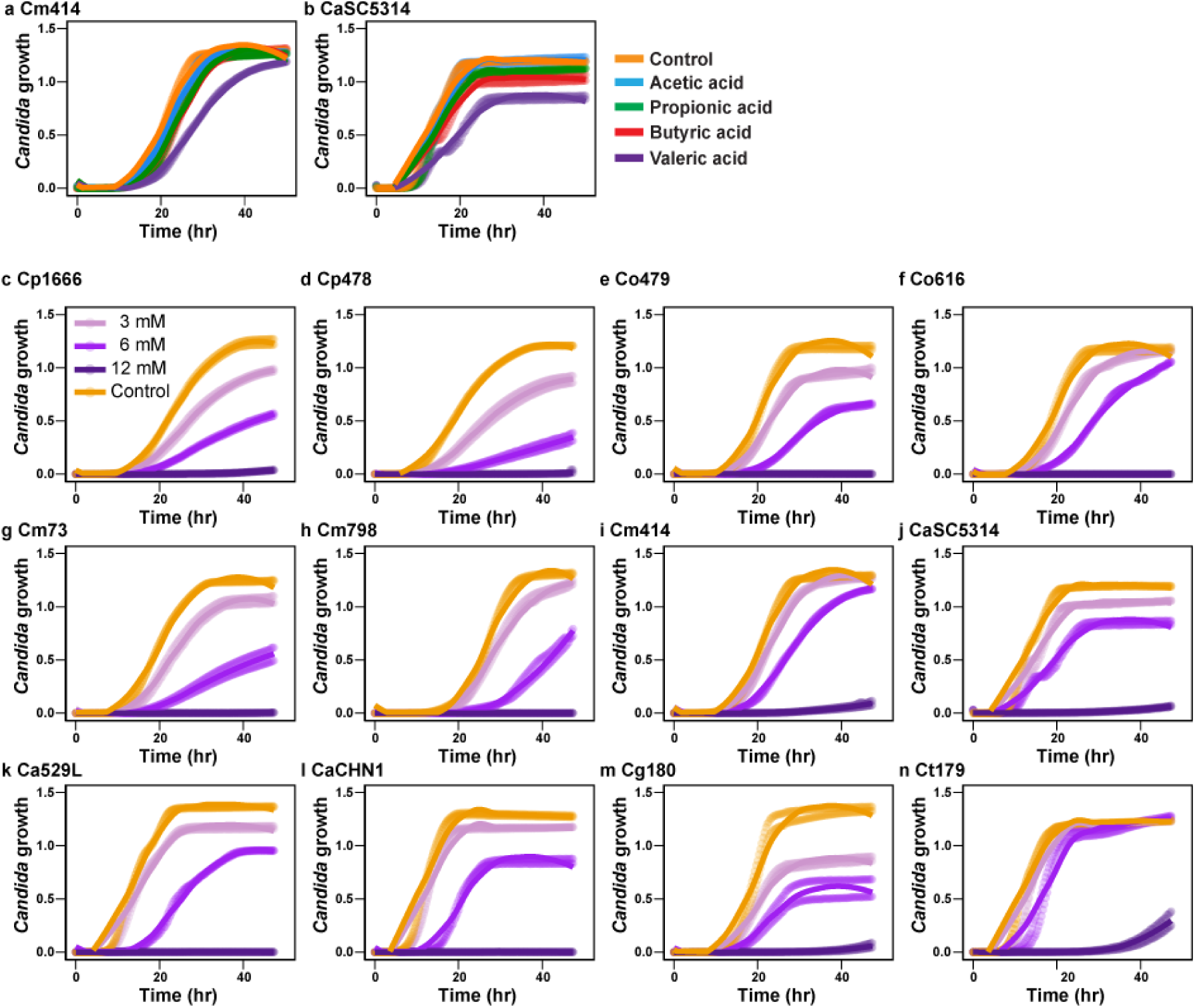
*C. parapsilosis* and other *Candida* species growth curves in the presence of valeric acid or other SCFAs, related to Figure 3. **a, b,** (**a**) *Candida metapsilosis* 414 and (**b**) *C. albicans* SC5314 growth by OD_600_ measurement in the presence of 0 mM SCFA (control, orange line) or 6 mM acetic acid (blue line), propionic (green line), butyric (red line), or valeric acid (purple line). **c-n**, *Candida* growth in 0 mM (control, orange), 3 mM (light purple), 6 mM (purple), or 12 mM (dark blue) valeric acid. (**c, d**) *C. parapsilosis* 1666 and 478, (**e, f**) *C. orthopsilosis* 479 and 616, (**g-i**) *C. metapsilosis* 73, 798, and 414, (**j-l**) *C. albicans* SC5314, 529L, and CHN1, (**m**) *C. glabrata* 180, and (**n**) *C. tropicalis* 179. Growth curves were calculated from at least two replicates per experiment.

**Extended Figure 5.**
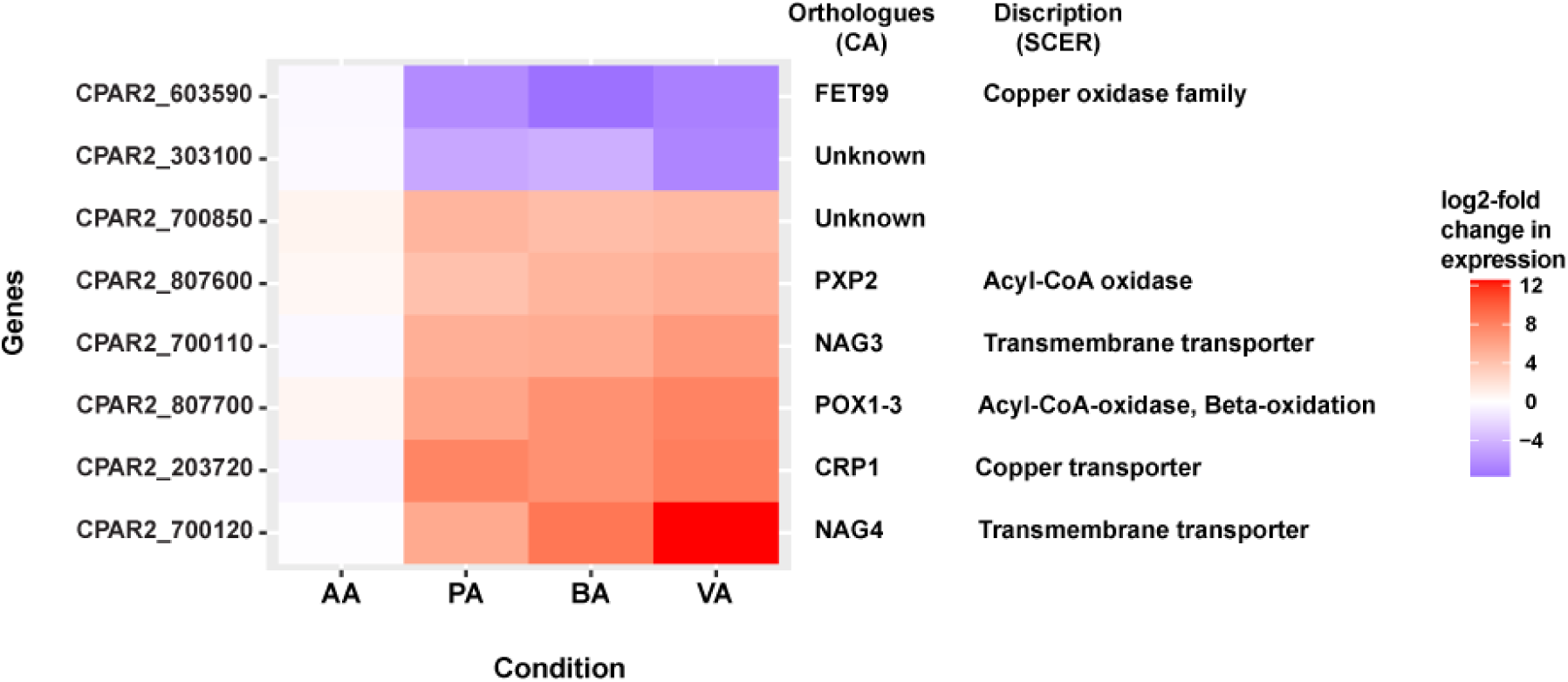
Differentially expressed genes in SCFA-treated *C. parapsilosis* samples, related to Figure 4. The heatmap depicts a subset of highly differentially expressed genes and annotations in propionic acid (PA)-, butyric acid (BA)-, and valeric acid (VA)-treated *C. parapsilosis* cells compared to untreated control or acetic acid (AA)-treated cells. *C. albicans* and *S. cerevisiae* orthologs and known gene functions are indicated in the two right columns. The heatmap includes genes with a log_2_-fold change > 4 or < −4 and an adjusted *P* value < 0.01 for PA-, BA- and VA-treated cells, and a log_2_-fold change < 1 or > −1 for AA-treated cells.

**Extended Figure 6.**
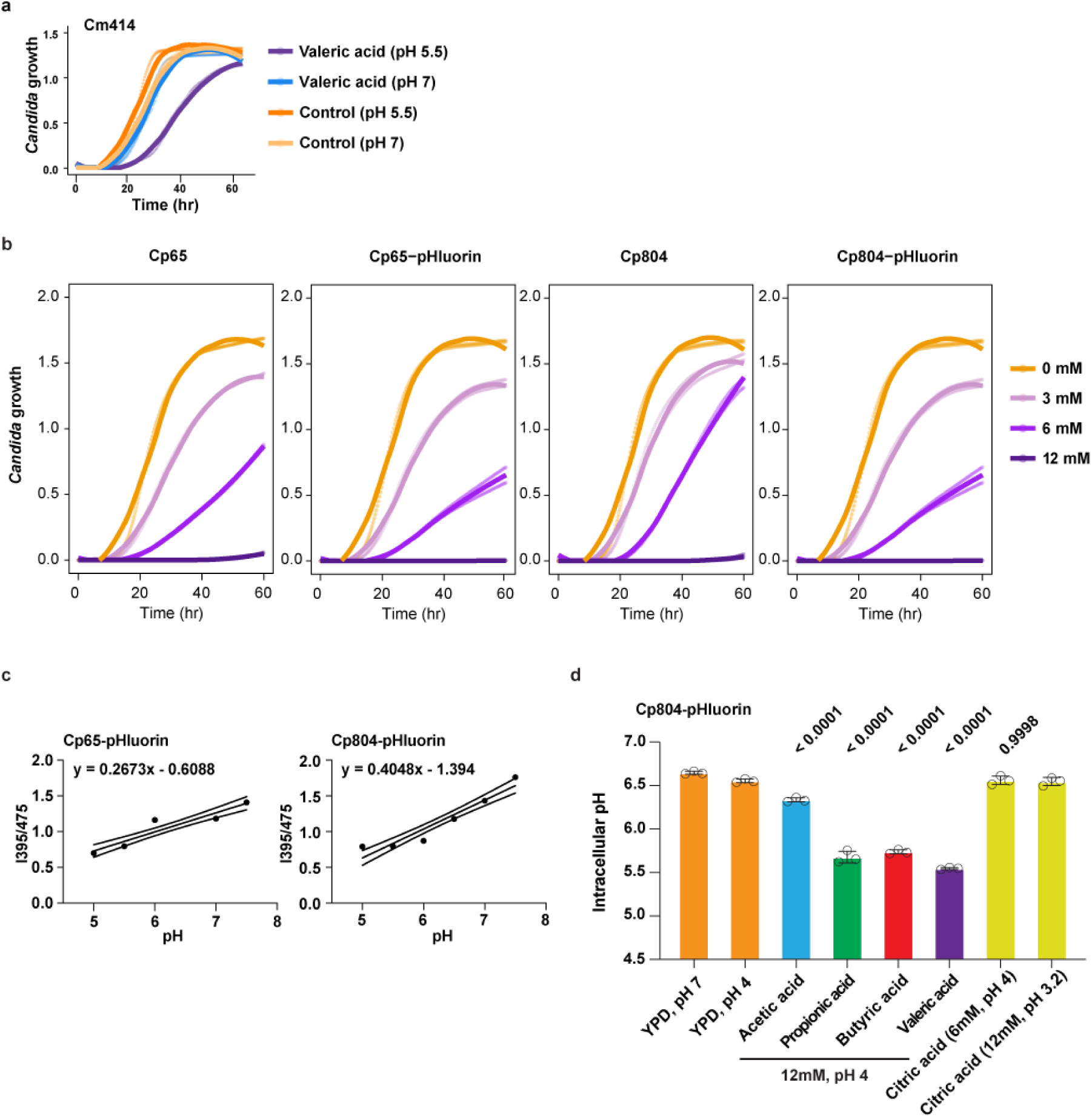
Characterization of pHluorin-expressing *C. parapsilosis* strains, related to Figure 5. **a,** *C. metapsilosis* strain 414 growth in 0 mM (pH 5.5, orange or pH 7, yellow) or in 6 mM valeric acid (pH 5.5, purple line or pH 7, blue line). **b**, Growth curves of *C. parapsilosis* 65 (Cp65), pHluorin-expressing *C. parapsilosis* 65 (Cp65-pHluorin), *C. parapsilosis* 804 (Cp804), pHluorin-expressing *C. parapsilosis* 804 (Cp804-pHluorin) in 0 mM (orange), 3 mM (light purple), 6 mM (purple), and 12 mM valeric acid (dark blue). The lines show the average OD_600_ values of duplicate samples. **c,** Standard curves of monensin-permeabilized Cp65-pHluorin and Cp804-pHluorin cells that indicate the 395nm/475nm emission ratios (y-axis) at the indicated pH (x-axis). **d,** Intracellular pH of Cp804-pHluorin cells after incubation in YPD broth with the indicated concentrations (0-12 mM, 0 mM control, orange) of acetic acid (light blue), propionic acid (green), butyric acid (red), valeric acid (purple), or citric acid (yellow), as determined by fluorescence emission quantitation at 509 nm with 395 and 475 nm excitation wavelengths using a plate reader. The intracellular pH was calculated using a standard curve of monensin-permeabilized pHluorin-expressing cells as in (**c**). Representative data from two independent experiments shown. Statistical analysis: **d**, Data as mean values are shown with adjusted *P* values versus pH-adjusted control or indicated pairs (*P* value: 12 mM of indicated short-chain fatty acid vs control pH 4, or 6 mM of citric acid vs control pH 4) calculated with 2way ANOVA with Bonferroni’s multiple comparison test.

**Extended Figure 7.**
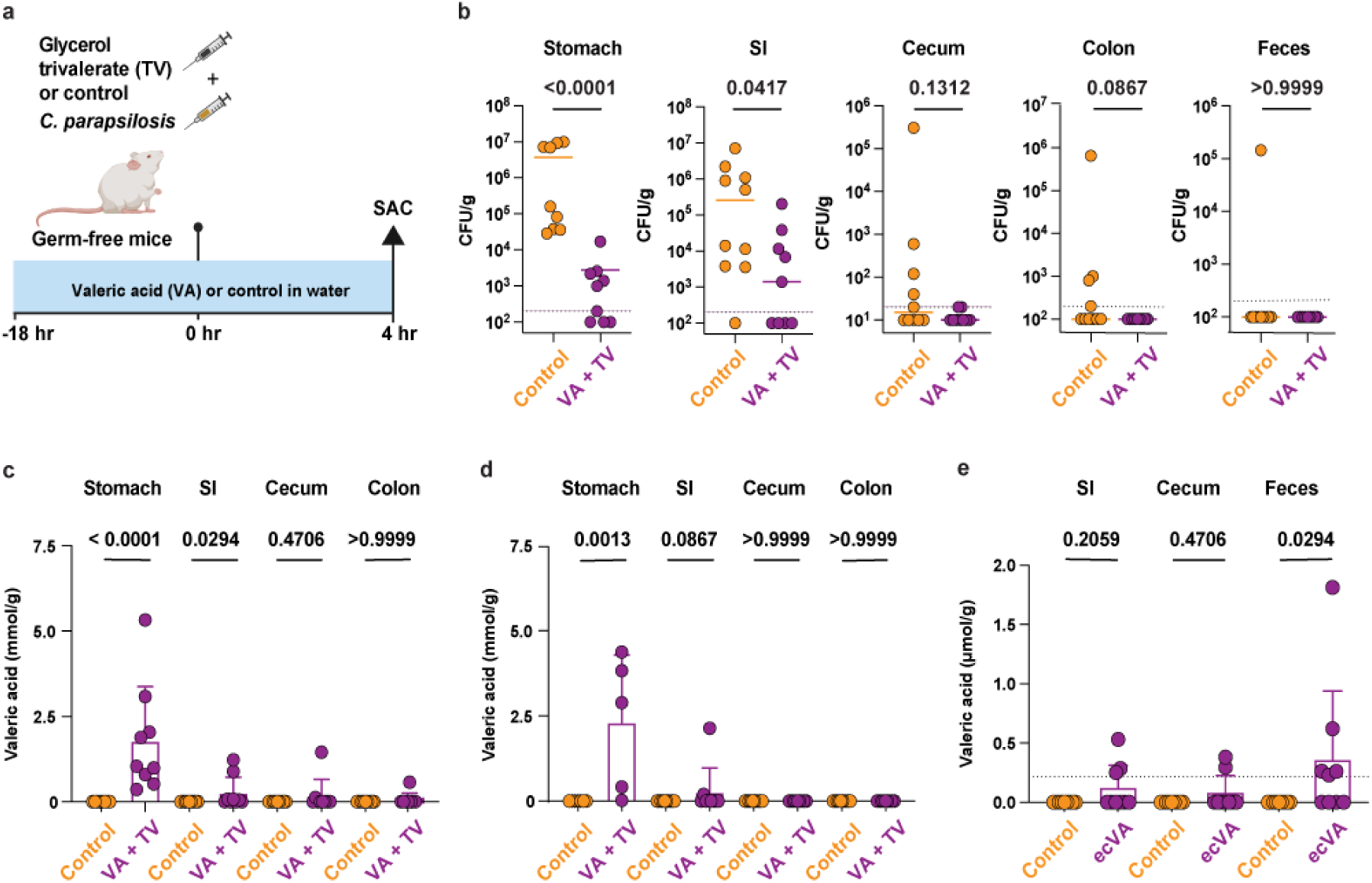
Valeric acid-associated inhibition of *C. parapsilosis* growth and valeric acid abundance in the intestinal tract, related to Figure 6. **a**, Experimental scheme to test the administration of glycerol trivalerate (TV, gavage), and valeric acid (VA, drinking water *ad libitum*) on *C. parapsilosis* intestinal colonization in germ-free Balb/c mice. **b**, *C. parapsilosis* CFU in stomach, small intestine (SI), cecal, or colonic contents, or in the feces of germ-free Balb/c mice that received 200 µl of 1M TV (purple dots) or equivalent glycerol (control, orange dots) and 1 x 10^6^ *C. parapsilosis* by gavage, with mice maintained on 0 mM (control group treated with glycerol) or 150 mM VA in the drinking water (TV group). **c-e**, Valeric acid levels in indicated intestinal contents from experiments shown in (**c**) Figure 6d, (**d**) Extended Figure 7b, and (**e**) Figure 6g. Results were pooled from two independent with 7-11 mice per group. Each dots represents one mouse. Statistical analysis: **b-e**, *P* values were calculated using two-tailed Mann–Whitney U test.

